# Calorie restriction intervention induces enterotype-associated BMI loss in nonobese individuals

**DOI:** 10.1101/514596

**Authors:** Hua Zou, Dan Wang, Huahui Ren, Peishan Chen, Chao Fang, Zhun Shi, Pengfan Zhang, Jian Wang, Huanming Yang, Kaiye Cai, Huanzi Zhong

## Abstract

Calorie restriction (CR), which has the potential effect to weight loss and blood amino acids, has been demonstrated to associate with gut microbiota in human, especially in obese individuals. However, studies for simultaneously evaluating enterotype-dependent impacts of CR on the gut microbiota and blood amino acids in nonobese individuals are still limited.

Here, 41 nonobese individuals received a 3-week CR diet with approximately 50% fewer calories than normal diet. We measured their BMI and blood amino acid concentration, along with the gut microbiota before and after the intervention. In this trial, 28 Enterotype *Bacteroides* (ETB) subjects and 13 Enterotype *Prevotella* (ETP) subjects were identified before the intervention. Short-term CR dietary intervention decreased the body mass index (BMI) in most subjects but varied in subjects with different enterotypes. ETP subjects exhibited significantly higher BMI loss ratio than the ETB subjects. CR additionally induced substantial enterotype-independent changes in blood amino acids, but only minor changes in gut microbial composition.

We further built a prediction model based on baseline relative abundances of 7 gut microbial species showing high performance in predicting CR-associated BMI loss ratio. Among them, the relative abundance of ETB-enriched *Clostridium bolteae* and *C. ramosum* were negatively correlated with BMI loss ratio while the relative abundance of *Dorea longicatena* which was slightly enriched in ETP subjects, was positively correlated with BMI loss ratio.

Together, our work points out that the individual variation of BMI loss after CR could be partially correlated with different microbial composition and highlights the potential application for microbiome stratification in personalized nutrition intervention.

## Introduction

Calorie restriction, a nutritional intervention of reduced energy intake, has been demonstrated to reduce body weight and modulate serum metabolic in human population in many studies [1, 2]. Observational studies with long-term CR found that CR resulted in weight loss and decreasing chronic disease risk factors in nonobese persons [3–6]. For instance, in a 2-year nonobese human trail, CR significantly decreased body weight and cardiometabolic risk factors, such as triglycerides and total cholesterol [4]. Another study [5] also provided evidence for reduced BMI and improved mood and sleep duration with CR in healthy nonobese adults. However, studies about potential negative impacts of CR on human health remain limited.

CR intervention affecting gut microbiota and blood amino acids has been explored in many studies, particularly among overweight and obese individuals [7–10]. A 4-week CR intervention improved gut barrier integrity, reduced systemic inflammation on gut microbial diversity and BMI loss in obese women, suggesting a potential association among gut microbiota, CR and BMI [8]. Increasing studies have demonstrated that the gut microbiota has been implicated in modulating host energy and nutrient metabolism [11] and regulating the production of gastrointestinal hormones and host appetite [12, 13]. Enterotype, a concept for stratifying individuals based on the gut microbiota, was first described in 2011 and was closely linked to long-term dietary patterns [14, 15]. Plenty of studies have reported that individuals have shown microbial-dependent (enterotypes, *Bacteroides* to *Prevotella* ratio) metabolic responses to the same intervention, including changes of BMI and glycemic indices [12, 15–20]. As for the association between CR and blood amino acids, Biolo *et.al* reported changes in blood amino acids of 9 healthy nonobese men after 2-week CR diet [10]. Despite the plenty of studies on CR as we mentioned above, to our knowledge, there are no studies for simultaneously evaluating impacts of CR on BMI, gut microbiota and blood amino acids in nonobese individuals in a single study.

Here, we conducted a 3-week CR intervention on 41 nonobese subjects with two enterotypes, including enterotype *Bacteroides* (ETB) and enterotype *Prevotella* (ETP). We found that ETP subjects had higher BMI loss ratio than ETB subjects after CR intervention. On the other hand, we found that there were no obvious changes in the gut microbiota composition of two enterotype groups after the CR intervention and the two enterotype groups showed consistent changes in levels of multiple blood amino acids. We further demonstrated that baseline relative abundances of 7 gut microbial species including 3 enterotype-specific species *(C. bolteae, C. ramosum,* and *D. longicatena),* rather than baseline BMI and blood amino acids levels, could well predict CR-associated BMI loss ratio. Additionally, we found significantly increased levels of 3-methylhistidine after the intervention, a potential biomarker for muscle protein turnover, suggesting possible muscle protein loss under a CR diet with lower than normal protein availability.

## Materials and methods

### Volunteer Recruitment

Volunteer-wanted posters were propagated at the China National Gene Bank in Shenzhen from March to April 2017. A non-obese healthy volunteer was considered if his/her BMI was less than 28 kg/m^2^ [21]. In addition, recruited volunteers should meet all the following criteria: 1) without antibiotics in the recent 2 months; 2) without prebiotic or probiotic supplements in the recent 2 months; 3) without hypertension, diabetes mellitus, gastrointestinal disease and other severe auto-immune disease; 4) regular eating and lifestyle patterns; 5) no international travel in the recent 3 month. 50 individuals met all the criteria and were recruited in this study, and 41 individuals (24 females and 17 males aged 30 ± 6 years old) completed the whole intervention (**Table 1**). The study was approved by the institutional review board on bioethics and biosafety of BGI-Shenzhen, Shenzhen (NO. BGI-IRB 17020). All participates were fully informed of the design and purpose of this intervention study and signed a written informed consent letter.

**Table 1.** Cohort description

|  | **Cohort**<br>**(Mean ± SD)** |
| --- | --- |
| **Number of subjects** | 41 |
| **Sex (female/male)** | 24/17 |
| **Age** | 30 ± 6 |
| **BMI (kg/m<sup>2</sup>)** | 23.72 ± 2.81 |
| **Weight(kg)** | 64.84 ± 11.57 |

### Study design and low-calorie food preparation

The study included a one-week run-in period (baseline) and a three-week CR dietary intervention period. During the first week (run-in period), all healthy volunteers consumed their usual diet and were encouraged to avoid yoghurt, high-fat foods and alcohol. The CR diet was comprised of ~50% calories of a normal-calorie diet (female, 1000kcal/day; male, 1200kcal/day). It was designed with carbohydrate, fat and protein as approximately 55%, 30% and 15% of the total energy intake respectively, according to the Dietary Guidelines for Chinese Residents (2016) and nutritionally balanced [22] and a recent large nutritional study in China [23]. Common foods in low-calorie diets such as rice, vegetables, eggs, pork and beef were prepared in our study center to control experimental variables introduced by different foods and calorie estimation errors.

Traditional Chinese cooking style - boiled, stir-fried and stewed, were applied for our foods. For each meal, digital scales were used to measure the nutritional and caloric values of different foods and total meal for male and female respectively.

BMI, blood and fecal samples of each volunteer were collected at our study center at baseline and after the 3-week CR intervention (**Figure 1**). To avoid intra-individual variations, BMI was collected multiple times of each volunteer during the last week of the CR intervention, and the averaged BMI value was used as his/her after-intervention BMI (**Figure 1**).

**Figure 1.**
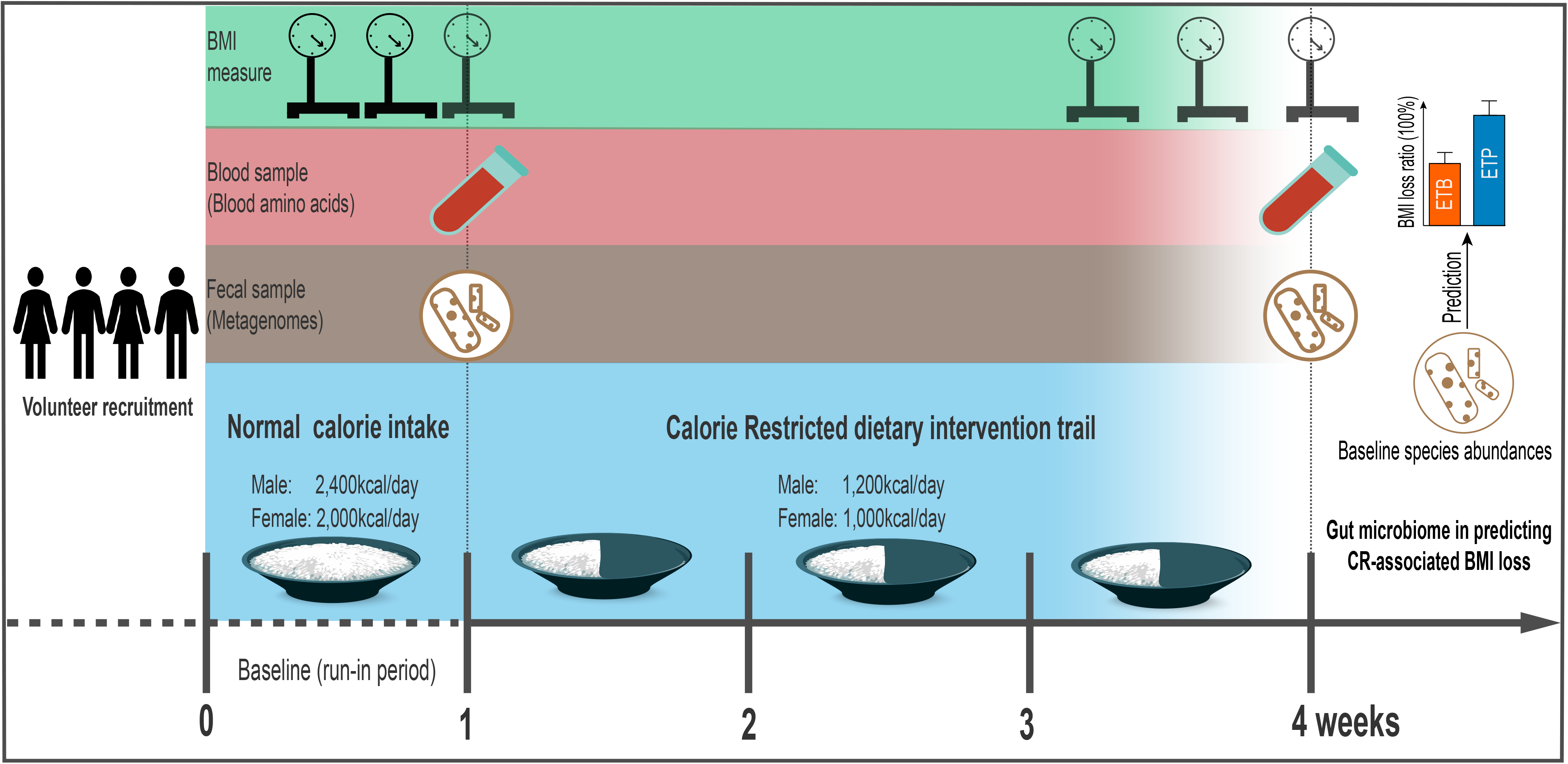
Overview of the experimental design. Illustration of our experimental design, including a 1-week run-in period (baseline) and a 3-week calorie restriction (CR) dietary intervention trial with 50% energy deficit diet (male, ~1200Kcal/day; female, ~1000Kcal/day). BMI, fasting blood samples and fecal samples of 41 enrolled healthy subjects were collected before and after the CR intervention to assess its effects on BMI, blood amino acids and gut microbiome in two enterotype groups.

### Fecal sampling and shotgun metagenomic sequencing

Fecal samples were self-collected and then transferred to the laboratory on dry ice and kept frozen at −80°C before and after the CR intervention. Fecal DNA was extracted following a manual protocol as described previously [24]. The DNA concentration was estimated by Qubit (Invitrogen). Library construction and shotgun metagenomics sequencing were performed on qualified DNA samples based on the BGISEQ-500 protocol in the single-end 100bp mode [25].

### Metagenomic analysis

Raw reads of BGISEQ-500 with SE100 mode were trimmed by an overall accuracy (OA) control strategy to control quality [25]. After trimming, averagely, 98.15% of the raw reads still remained as high-quality reads (**Supplemental Table 1**). By using SOAP2.22 software, the high-quality reads were aligned to hg19 to remove reads from host DNA (identity ≥ 0.9). The retained clean reads were aligned to the integrated non-redundant gene catalog (IGC) using SOAP2.22 [26] and the average mapping rate and unique mapping rate were 80.18% and 65.76% respectively (identity ≥ 0.95, **Supplemental Table 1**). The relative abundance profiles of genes, genera, species and Kyoto Encyclopedia of Genes and Genomes orthologous groups (KEGG, KOs) of each sample were calculated by summing the relative abundances of their assigned IGC genes [26].

For enterotyping, we applied a recently published universal classifier (http://enterotypes.org/), which circumvents major shortcomings in enterotyping methodology such as lack of standard and small sample size [27].

At baseline, 41 individuals were clustered into two groups: 28 ETB *(Bacteroides* enriched) and 13 ETP *(Prevotella* enriched) individuals 87.8% (36 of 41) individuals were clustered to the same enterotype after the 3-week CR intervention. Detailed enterotype information for each individual is provided in **Supplemental Table 2**.

Genus or species with an occurrence rate > 80% and a median relative abundance > 1e-6 in all samples were defined as common genus or species and used for further intra- and inter-enterotype comparison analyses **Supplemental Table 3-4**.

Differentially enriched KEGG pathways were identified between enterotypes and between different time points, based on the distribution of Z-scores of all KOs belonging to a given pathway [28, 29]. A reporter score |Z| > 1.96 (95% confidence interval according to a normal distribution) was used as a detection threshold for significantly differentiating pathways.

Alpha diversity of each individual was calculated on the gene and species relative abundance profiles using the Shannon index. Beta-diversity on the gene and species relative abundance profiles was calculated using the Bray-Curtis distance.

### Blood sample collection and amino acids profiling

Fasting blood samples were collected before and after the intervention for amino acids analysis. These blood samples were then centrifuged, and serum samples were collected and stored at −80 _. The concentrations of 31 amino acids and derivatives in the serum samples were then measured via ultra-high pressure liquid chromatography (UHPLC) coupled to an AB Sciex Qtrap 5500 mass spectrometry (AB Sciex, US) as described previously [30].

### Statistical methods

#### Pearson’s chi-square test

Pearson’s chi-square test was performed to assess sex distribution between individuals of two enterotypes.

#### Wilcoxon rank-sum test & Wilcoxon Signed-rank test

Wilcoxon rank-sum test was used to detect the significant differences on phenotypes, the concentrations of blood amino acids and the relative abundances of genera and species between enterotypes.

Wilcoxon signed-rank test was used to detect the significant differences on phenotypes, the concentrations of blood amino acids and the relative abundances of genera and species in paired samples before and after the intervention.

#### BMI loss ratio

BMI loss ratio of a given individual was calculated using the following equation:

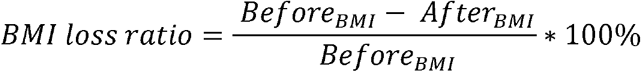

Where *Before_BMI_* and *After_BMI_* are the BMI value of the same individual before and after the CR intervention, respectively.

## PERMANOVA

The association between enterotypes and the overall blood amino acid profile at baseline was assessed using permutational multivariate analysis of variance (PERMANOVA) with 9,999 permutations on enterotypes (R *vegan* package, *adonis* function, method=“bray”).

### PCoA

Principal coordinate analysis (PCoA) of fecal samples was performed based on the relative abundances of common species using Bray-Curtis distance (R *ape* package).

### PCA

Principal component analysis (PCA) was performed based on the blood amino acid profiles to visual overall amino acid composition between enterotypes, and that before and after the intervention.

### Feature selection of gut species and serum amino acids

To investigate whether we could predict BMI loss ratio using omics features, we performed a Lasso (Least absolute shrinkage and selection operator) regression analysis between baseline relative abundances of gut common species and the concentrations of blood amino acids (independent variables), and BMI loss ratio (dependent variables).

We first normalized values of both independent and dependent variables (R, *scale* function). We then used the R function cv.glmnet to choose the most appropriate value for *λ* in the Lasso model (R *glmnet* package, alpha =1, family=“gaussian”, nfolds=10, alpha=1, nlambda=100). Here, *λ* is the tuning parameter (*λ*□>□0) which controls the strength of the shrinkage of the variables [31]. We then applied the Lasso feature selection process by shrinking the Lasso regression coefficients of non-informative variables to zero and selecting the variables of non-zero coefficients. Seven gut microbial species including *Clostridium bolteae, Clostridium ramosum Dorea longicatena, Coprococcus eutactus, Streptococcus mitis, Clostridiales genomosp. BVAB3* and *Mobiluncus curtisii* were selected at this step.

### Performance estimation of BMI loss ratio prediction model

To reduce overfitting with a limited sample size (n=41), we applied leave-one-out cross validation (LOOCV) to estimate the prediction performance of BMI loss ratio using a generalized linear model (GLM) of the seven selected features (creatFolds function in R *caret* package and the glm function in R *base* package). Likewise, we also used baseline BMI values for LOOCV to estimate its prediction performance for CR-associated BMI loss ratio. Spearman’s rho values were calculated between actual BMI loss ratios and the predicted values.

### Significance cutoff

P-value adjustment was applied for multiple hypothesis testing on the concentrations of blood amino acids, the relative abundances of gut microbial genera and species used Benjamini-Hochberg (BH) method. BH-adjusted P value less than 0.05 was considered as statistical significance. The significance for α-diversity, β-diversity and phenotypes (age, female to male ratio, BMI and BMI loss ratio) was set at p < 0.05.

All statistical analyses were conducted using R (version 3.5.0).

## Result

### BMI loss of ETB and ETP subjects responded differentially to CR Intervention

Based on the baseline genera abundance profile, individuals can be robustly clustered into two enterotypes: enterotype *Bacteroides* (ETB, n = 28) and enterotype *Prevotella* (ETP, n = 13) (See Materials and Methods, **Figure 2A**).

**Figure 2.**
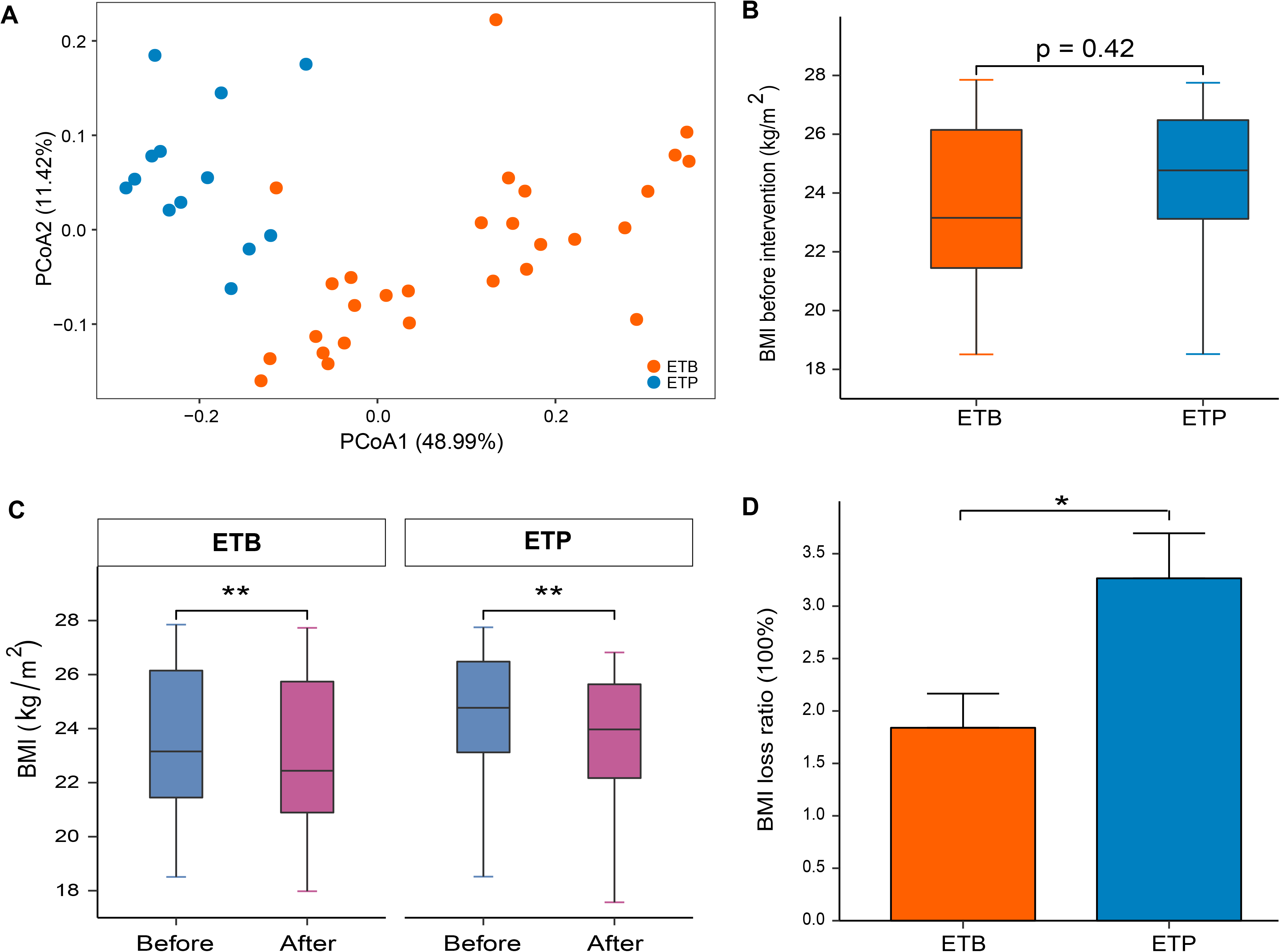
A short-term CR intervention altered BMI. **(A)** Principal coordinates analysis (PCoA) based on genera-level Bray-Curtis distance between all baseline fecal samples. Orange, subjects of enterotype *Bacteroides* (ETB) and blue, subjects of enterotype *Prevotella* (ETP). **(B)** Baseline BMI between ETB and ETP subjects. **(C)** Changes in BMI before and after intervention in individuals of each enterotype. (D) Boxplot showing BMI loss ratio between ETB subjects and ETP subjects. *, P < 0.05; **, P < 0.01.

Comparisons of the baseline phenotypes between two enterotype groups revealed that all collected phenotypes, including sex distribution (female/male ratio), age, BMI and weight, showed no significant differences between ETB and ETP subjects (**Table 2, Figure 2B**). By contrast, the baseline compositional and functional characteristics of the gut microbiota showed marked differences between two groups, in agreement with the previous studies [12, 15, 32]. For instance, genera *Prevotella* and *Paraprevotella* and four species from the two genera were significantly enriched in ETP subjects whereas 19 common species, from such as genera *Bacteroides* and *Clostridium* including *C. bolteae* and *C. ramosum,* were significantly enriched in ETB subjects (Wilcoxon rank-sum test, BH-adjusted P < 0.05; **Supplemental Figure 1A, Supplemental Table 5-6**). At the functional level, multiple pathways in carbohydrate metabolism and membrane transport were highly enriched in ETB subjects whereas four pathways including biosynthesis of phenylalanine, tyrosine and tryptophan (map00400), peptidoglycan (map00550), terpenoid backbone (map00900); and methane metabolism (map00680) were enriched in ETP subjects (|Z-score| > 1.96, **Supplemental Figure 1B, Supplemental Table 7**).

**Table 2.** Comparison of baseline phenotypes between ETB and ETP subjects

|  | **ETB Group**<br>**(Mean ± SD)** | **ETP Group**<br>**(Mean ± SD)** | **P value**<br>**ETB vs ETP** |
| --- | --- | --- | --- |
| **Number of subjects** | 28 | 13 |  |
| **Sex (female/male)** | 16/12 | 8/5 | 1 |
| **Age** | 29 ± 6 | 30 ± 7 | 0.44 |
| **BMI (kg/m<sup>2</sup>)** | 23.50 ± 2.81 | 24.21 ± 2.85 | 0.42 |
| **Weight (kg)** | 64.40 ± 11.37 | 65.81 ± 12.39 | 0.62 |

After the 3-week CR intervention, BMI values were decreased significantly in both ETB and ETP subjects (**Figure 2C**; Wilcoxon Signed-rank test, p < 0.05). Interestingly, subsequent analysis revealed that ETP subjects showed significantly greater BMI loss ratio than ETB subjects (Wilcoxon rank-sum test, p < 0.05; mean BMI loss ratio 3.27% versus 1.84%; **Figure 2D**; see Materials and Methods).

### Overall gut microbiome composition is stable to CR Intervention

We next investigated the extent and impacts of the CR intervention on gut microbial composition in subjects of different enterotypes. Principal coordinate analysis (PCoA) based on species abundance profile of all samples showed that the projected coordinates of each enterotype group did not change significantly before and after the intervention (**Figure 3A**, p > 0.05). Furthermore, 23 of 28 ETB subjects and 12 of 13 ETP subjects were assigned to the same enterotypes after the intervention (**Supplemental Table 2**). In addition, α-diversity (**Figure 3B-C**) and β-diversity (**Figure 3D-E**) at the genera and species level of fecal samples also showed no significant changes before and after the intervention in two enterotype groups, respectively (Wilcoxon Signed-rank test, P > 0. 05). We further revealed that no common species differed significantly in abundance before and after the intervention in each enterotype group (**Supplemental Table 8**, BH-adjusted P > 0.05). All these findings suggest overall stable gut microbial composition in response to a 3-week 50% energy deficit CR intervention.

**Figure 3.**
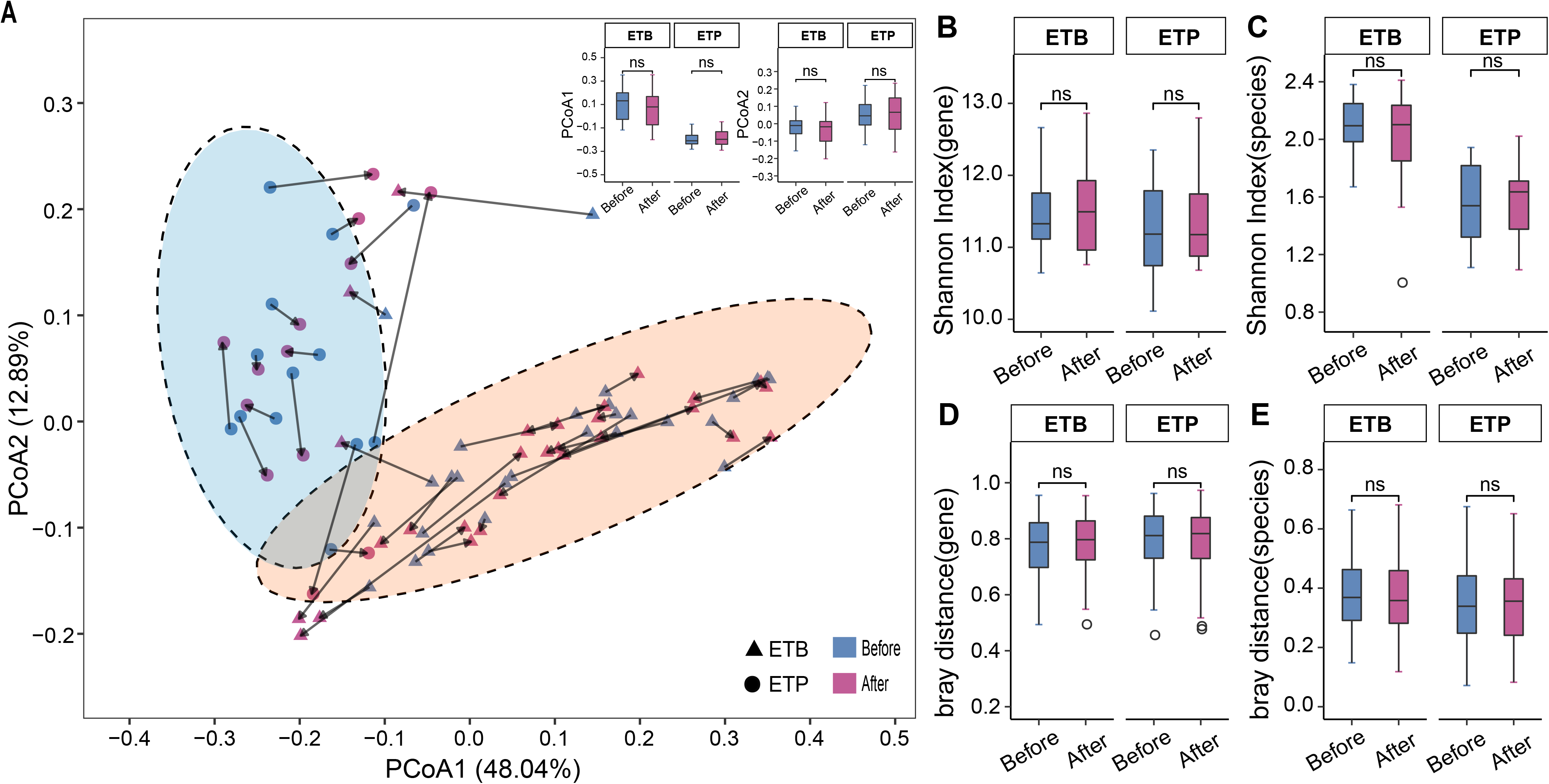
Overall gut microbial composition of two enterotypes before and after the CR. **(A)** Species-based principal coordinates analysis (PCoA) of subjects before and after the CR trial. Trangers, samples of ETB; Circles, samples of ETP. Arrows indicate paried samples from the same indivudul. Boxplot showing the projected coordinate 1 (PCo1) and PCo2 of samples before and after the intervention. **(B-C)** α-diversity (Shannon index) at the gene and species levels before (blue) and after (red) intervention in each enterotype group. **(D-E)** β-diversity (Bray-Curtis distance) at the gene and species levels before (blue) and after (red) the intervention in each enterotype group.-ns, no significance, P > 0.05, Wilcoxon signed-rank test.

On the other hand, we observed that the abundances of several functional pathways were changed after the intervention. For instance, abundances of pathways for pentose and glucuronate interconversions (map00040) and galactose metabolism (map00052) were significantly decreased after the CR intervention in both enterotype groups. ETB individuals additionally exhibited significantly reduced levels of pathways involved in cofactors metabolism (pantothenate and CoA biosynthesis, map00770; biotin metabolism, map00780) and increased levels of pathways for amino acids metabolism (phenylalanine metabolism, map00360; cysteine and methionine metabolism, map00270) after the intervention (**Supplemental Table 9**).

### Enterotype-independent alterations of blood amino acids to CR Intervention

Reflecting differential gut microbial functional potentials including amino acid metabolism between two enterotype groups, we asked whether blood amino acid composition was associated with enterotype. PCA of baseline amino acid profiles showed no separation of subjects of two enterotypes (**Figure 4A**, PERMANOVA, p > 0.05, See Material and Methods). In line with this result, we found no significant differences of baseline levels of blood amino acids between two enterotype groups (Wilcoxon rank-sum test, BH-adjusted P > 0.05, Figure 4B, Supplemental Table 10).

**Figure 4.**
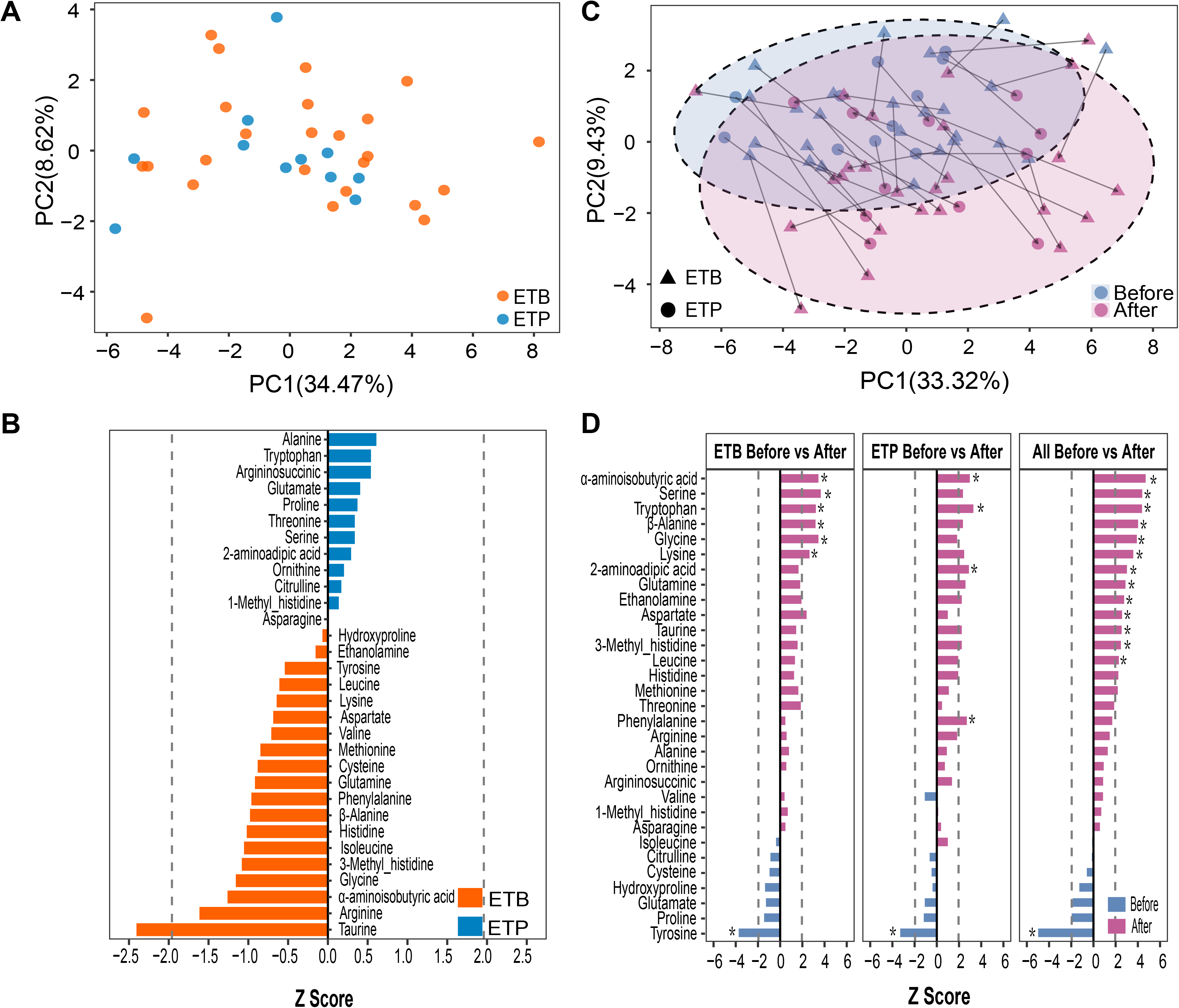
A short-term CR intervention altered blood amino acids. **(A)** Principal component analysis (PCA) of 41 subjects using baseline blood amino acid profiles. Orange, ETB; blue, ETP. **(B)** Comparison of baseline blood amino acid concentrations between ETB subjects (orange) and ETP subjects (blue). Wilcoxon rank-sum test, P values are transformed to Z-scores to represent enrichment directions. **(C)** Amino acid based PCA of samples before and after the intervention. Trangers, samples of ETB subjects; Circles, samples of ETP subjects. Arrows indicate paried samples from the same indivudul. **(D)** Changes in blood amino acid concentrations of ETB subjects, ETP subjects and all subjects before and after the intervention. Wilcoxon rank-sum test, P values are transformed to Z-scores to represent enrichment directions. Dashlied line indicates the absolute Z score of 1.96 (P = 0.05). Asterisk (*) indicates significance of Benjamini-Hochberg (BH) adjusted P < 0.05.

We next examined the potential impacts of the CR diet on blood amino acids. Notably, we observed similar changes in multiple blood amino acid concentrations in subjects of both enterotypes in response to the CR intervention (**Figure 4C, Supplemental Table 11**). We therefore combined all samples and found that levels of 13 blood amino acids or amino acid derivatives such as α-aminoisobutyric acid, β-alanine, glycine, serine, lysine and its degradation product 2-aminoadipic acid, and 3-methyl-histidine were significantly increased whereas only one measured amino acid tyrosine was significantly decreased after the intervention (Wilcoxon Signed-rank test, BH-adjusted P < 0.05, Figure 4D). In addition, we observed no significant differences in levels of blood amino acids between ETB and ETP subjects after the CR intervention (Wilcoxon rank-sum test, BH-adjusted P > 0.05, **Supplemental Table 12**), suggesting the effects of the CR intervention on blood amino acids were enterotype independent.

### Prediction of BMI loss ratio induced by CR intervention using gut microbial species

Considering the differential response in the BMI loss ratio to the CR intervention in two enterotypes, we next asked whether we could predict BMI loss ratio from the baseline omics measures. We thus built a Lasso shrinkage model between baseline levels of gut microbial species and blood amino acids and BMI loss ratio (**See Materials and Methods**). We successfully selected 7 gut microbial species showing associations with BMI loss ratio (Coefficient estimate > 0, **Figure 5A**). Interestingly, the relative abundances of 2 selected species *C. bolteae* and *C. ramosum,* which were enriched in ETB (Wilcoxon rank-sum test, BH-adjusted P < 0.05, **Supplemental Figure 1A, Supplemental Table 8**), were negatively correlated with BMI loss ratio. On the other hand, the relative abundance of *D. longicatena* which was slightly enriched in ETP (Wilcoxon rank-sum test, BH-adjusted P = 0.06) was positively correlated with BMI loss ratio (**Figure 5A**). Baseline abundances of the other 4 Lasso selected species, however, showed no significant differences between two enterotype groups (BH-adjusted P > 0.05, **Supplemental Table 6**). Among them, the abundances of *Coprococcus eutactus, Streptococcus mitis* and *Clostridiales genomosp. BVAB3* were positively associated with BMI loss ratio whereas the abundance of *Mobiluncus curtisii* was negatively associated with BMI loss ratio (**Figure 5A**).

**Figure 5.**
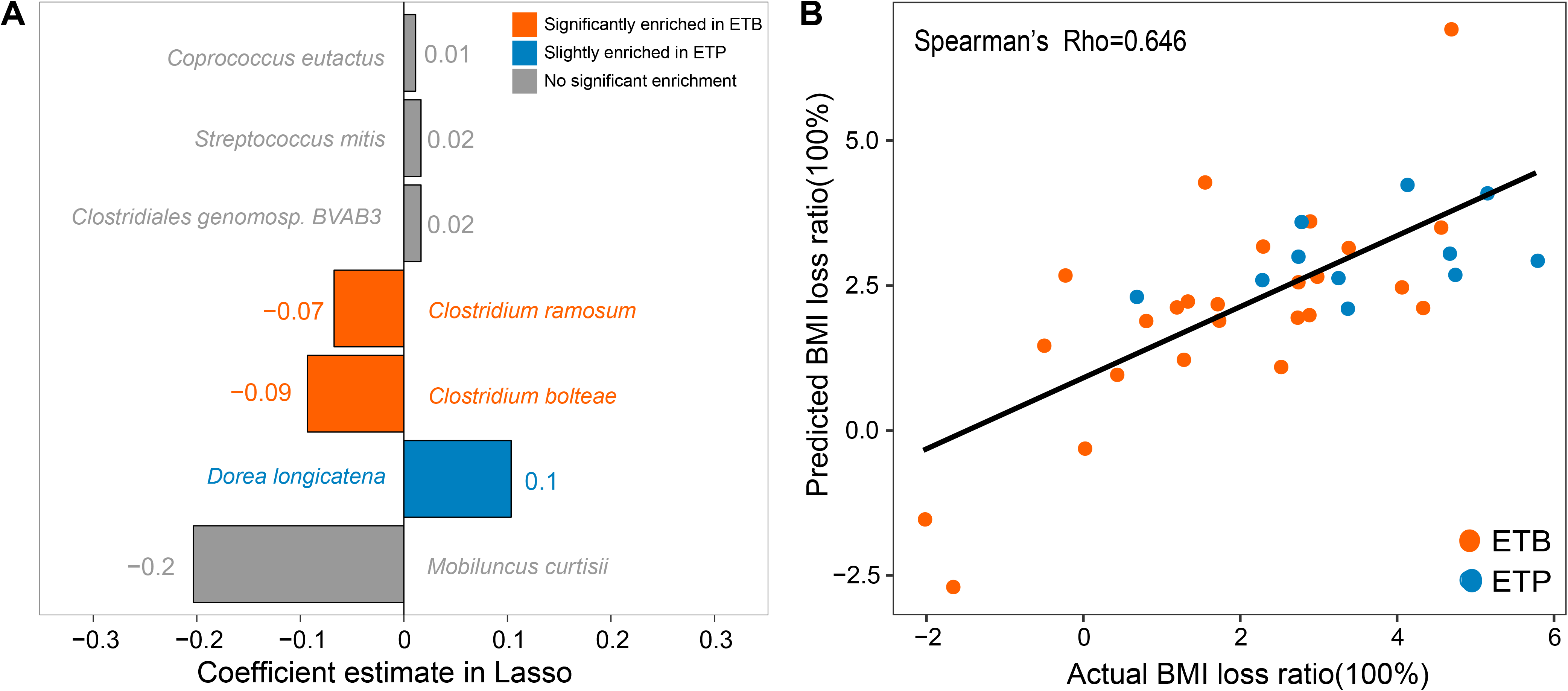
Prediction of BMI loss ratio using baseline abundances of gut microbial species. **(A)** Bar plot showing the 7 gut microbial species selected by least absolute shrinkage and selection operator (Lasso). Bar length indicates regression coefficient of each species estimated by Lasso. Orange, species significantly enriched in ETB subjects (BH-adjusted P < 0.05); blue, species slightly enriched in ETP subjects (P < 0.05 and BH-adjusted P = 0.06); grey, species with no significant enrichment between two enterotypes (P > 0.05). **(B)** Scatter plot showing prediction performance of BMI loss ratio based on the 7 selected species. Leave-one-out cross validation (LOOCV) was applied to evaluate the performance of generalized liner model (GLM), showing a strong Spearman’s rho between actual BMI loss ratios and predicted BMI loss ratios of 0.646. Red circles, ETB individuals; blue circles, ETP individuals.

To estimate the performance of 7 gut microbial species on the prediction of BMI loss ratio, we applied a general linear model between predicted BMI loss ratios and true values using leave-one-out cross validation (LOOCV). Notably, the result showed the predicted BMI loss ratio was significantly correlated with the true BMI loss ratio with a Spearman’s rho of 0.646 (**Figure 5B**, See Materials and Methods). By contrast, we found that the individual baseline BMI could hardly predict their BMI loss after the CR intervention (Spearman’s rho = −0.016, **Supplemental Figure 2**).

## Discussion

Here we evaluated changes in BMI, blood amino acids and the gut microbiota of 41 nonobese subjects stratified by enterotype after a 3-week 50% energy deficit CR intervention. We revealed that CR had significant effects on BMI and ETP subjects had higher BMI loss ratio than ETB subjects. Additionally, CR showed only minor impacts on gut microbial composition but significantly impacted blood amino acids in an enterotype-independent manner. We further selected 7 gut bacterial species whose baseline relative abundances could well predict BMI loss ratio induced by the CR intervention.

In this study, nonobese ETP individuals showed greater BMI loss than ETB individuals after our 3-week intervention. Similar findings were also reported in overweight individuals that high *Prevotella/Bacteroides* ratio group had higher weight loss than low *Prevotella/Bacteroides* ratio group when receiving the New Nordic Diet [17] and a 500 kcal/day energy deficit diet [9]. Importantly, we successfully constructed a BMI loss ratio prediction model based on baseline relative abundance of 7 gut microbial species. Among them, *C. ramosum* and *C. bolteae* (negatively associated with BMI loss ratio in our model) were significantly more enriched in ETB subjects while *D. longicatena* (positively associated with BMI loss ratio in our model) was slightly more enriched in ETP subjects. These three enterotype-specific species in our prediction model also have been reported by several other studies. A previous study showed that *C. ramosum* could enhance high-fat-diet-induced obesity in gnotobiotic mice, probably by enhancing nutrient absorption [33]. Two human studies have also reported a positive link between the abundance of *C. ramosum* and several obese-related metabolic parameters [34, 35]. In an obese postmenopausal women study, the fecal abundance of *C. bolteae* was positively correlated with multiple metabolic markers such as fasting glucose and insulin resistance, whereas *D. longicatena* was negatively correlated [36].

Combining the results of our prediction model and all the above-mentioned studies, the different enrichment of *C. ramosum, C. bolteae* and *D. longicatena* between ETB and ETP subjects may help explain why ETP subjects showed significantly larger BMI loss ratio than ETB subjects. Moreover, all selected 7 species annotated to genera *Coprococcus, Streptococcus, Clostridium, Dorea* and *Mobiluncus* were not the main contributors to enterotypes, suggesting the need for human gut microbial stratification in a higher resolution than enterotypes in further microbiome-based personalized intervention studies.

Attention should also be paid that levels of multiple amino acids and their derivatives were significantly increased in healthy non-obese subjects after the CR intervention, including 3-methylhistidine, a well-established biomarker of skeletal muscle protein turnover [37, 38]. Previous studies have reported that skeletal muscle acted as a fuel reserve to provide glucose disposal for other organs during fasting in human [38, 39]. Since the CR in our study was 50% of normal energy-intake and contained much lower protein than a normal diet, it can be speculated that skeletal muscle proteins might be partially broken down for supplying energy. By contrast, another previous study which provided a high-soy-protein diet induced weight loss without losing muscle mass in obese and pre-obese subjects [40]. All these findings suggest that the availability of adequate dietary proteins in further diet interventions is important to preserve healthy muscle mass during weight loss.

## Acknowledgements

We are very grateful to acknowledge colleagues from the China National Genebank for preparing calorie restriction diets, and for DNA extraction, library construction and sequencing.

## Contributions

Z.H.Z and K.Y.C designed the project. P.S.C and K.Y.C oversaw the collection of fecal and blood samples and phenotypic data. H.Z, D.W, H.H.R and Z.H.Z established the concept and analysis framework of the study. H.Z, H.H.R and D.W performed the bioinformatic analysis. H.Z wrote the first version of the manuscript. D.W and Z.H.Z substantially revised the manuscript. All authors contributed to and approved the final manuscript.

## Funding source and disclosure of conflicts of interests

The authors declare that they have no competing interests. This work was funded by National Key Research and Development Program of China (No.2017YFC0909703) and support of Shenzhen Municipal Government of China (No. CXB201108250098A).

## Ethics approval and consent to participate

The study protocol was approved by the institutional review board on bioethics and biosafety of BGI-Shenzhen, Shenzhen (NO. BGI-IRB 17020). Written informed consent was obtained from all participants before the CR intervention.

## Availability of data and material

The metagenomic sequencing data of this study have been deposited in the China National Genebank Nucleotide Sequence Archive (CNSA) (https://db.cngb.org/cnsa/) under the BioProject number CNP0000247. The scripts and gut microbial profiles used to perform statistical analysis and figure plotting are available at https://github.com/HuaZou/NonobeseBP.

## supplemental materials

**Supplemental Figure 1.**
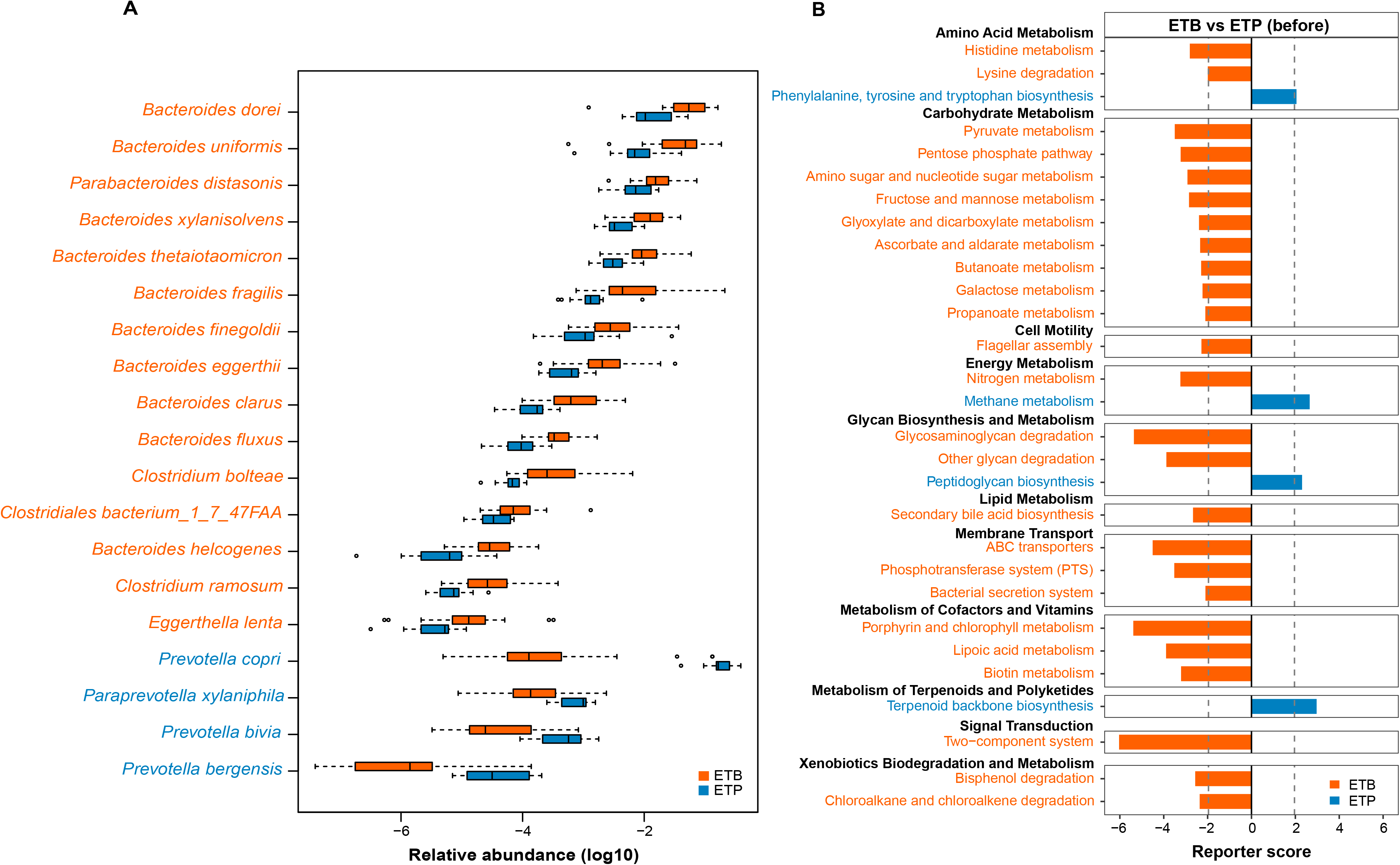
Baseline gut microbial differences between two enterotype groups. **(A)** Differentially enriched species between two enterotype groups. Orange and blue represent species overrepresented in ETB and ETP subjects, respectively. **(B)** Differential enrichment of KEGG pathways between ETB and ETP subjects. Dashed lines indicate a reporter score of 1.96, corresponding to 95% confidence in a normal distribution. Orange and blue bars indicate reporter scores of selected KEGG pathway overrepresented in ETB and ETP subjects, respectively.

**Supplemental Figure 2.**
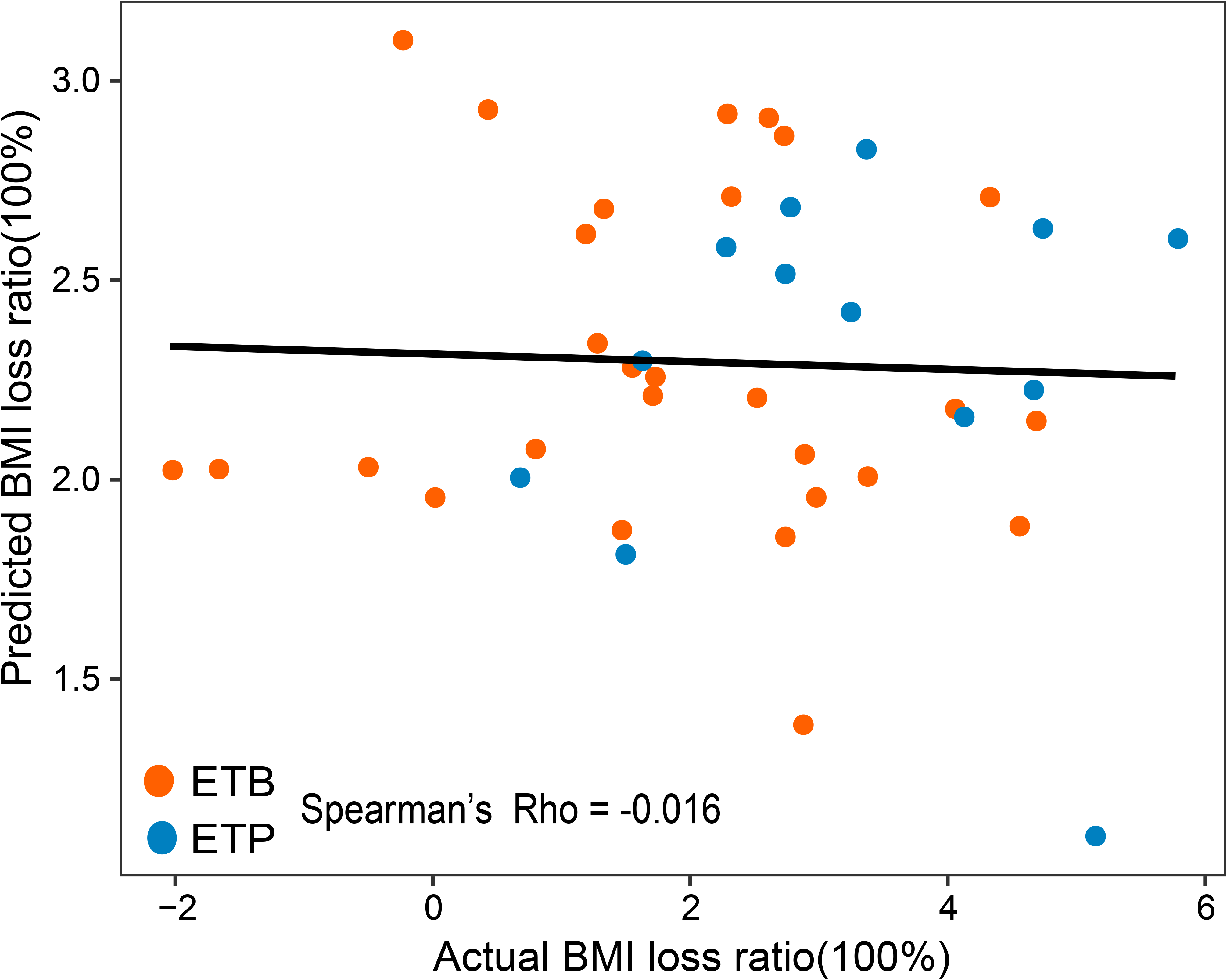
Performance of baseline BMI for prediction of BMI loss ratio. Scatter plot showing prediction performance of BMI loss ratio using baseline BMI values. A Spearman’s rho between actual BMI loss ratios and predicted BMI loss ratios was – 0.016. Red circles, ETB individuals; blue circles, ETP individuals.

**Supplemental Table 1.** Statistics for metagenomic sequencing data of fecal samples

**Supplemental Table 2.** Enterotype classification of individual fecal samples before and after the intervention

**Supplemental Table 3.** List of common genera

**Supplemental Table 4.** List of common species

**Supplemental Table 5.** Comparison of baseline genera relative abundance between ETB and ETP subjects

**Supplemental Table 6.** Comparison of baseline species relative abundance between ETB and ETP subjects

**Supplemental Table 7.** Differential enrichment of KEGG pathway between ETB and ETP subjects at baseline

**Supplemental Table 8.** Comparison of species relative abundance before and after the intervention in each enterotype

**Supplemental Table 9.** Differential enrichment of KEGG pathway before and after the intervention in each enterotype

**Supplemental Table 10.** Comparison of baseline blood amino acid levels between ETB and ETP subjects

**Supplemental Table 11.** Comparison of blood amino acid levels before and after the intervention

**Supplemental Table 12.** Comparison of blood amino acid levels between ETB and ETP subjects after the intervention

## Notes

#### Summary of Updates

Adjusted the orders of the table

